# Cohesin components Stag1 and Stag2 differentially influence haematopoietic mesoderm development in zebrafish embryos

**DOI:** 10.1101/2020.10.19.346122

**Authors:** Sarada Ketharnathan, Anastasia Labudina, Julia A. Horsfield

**Affiliations:** University of Otago, Department of Pathology, Otago Medical School, Dunedin, New Zealand; CHEO Research Institute, University of Ottawa, Ottawa, Canada; The University of Auckland, Maurice Wilkins Centre for Molecular Biodiscovery, Private Bag 92019, Auckland, New Zealand

**Keywords:** zebrafish, cohesin, haematopoiesis, mesoderm, development

## Abstract

Cohesin is a multiprotein complex made up of core subunits Smc1, Smc3 and Rad21, and either Stag1 or Stag2. Normal haematopoietic development relies on crucial functions of cohesin in cell division and regulation of gene expression via three-dimensional chromatin organisation. Cohesin subunit STAG2 is frequently mutated in myeloid malignancies, but the individual contributions of Stag variants to haematopoiesis or malignancy are not fully understood. Zebrafish have four Stag paralogues (Stag1a, Stag1b, Stag2a and Stag2b), allowing detailed genetic dissection of the contribution of Stag1-cohesin and Stag2-cohesin to development. Here we characterize for the first time the expression patterns and functions of zebrafish *stag* genes during embryogenesis. Using loss-of-function CRISPR-Cas9 zebrafish mutants, we show that *stag1a* and *stag2b* contribute to primitive embryonic haematopoiesis. Both *stag1a* and *stag2b* mutants present with erythropenia by 24 hours post-fertilisation. Homozygous loss of either paralog alters the number of haematopoietic/vascular progenitors in the lateral plate mesoderm. The lateral plate mesoderm zone of *scl*-positive cells is expanded in *stag1a* mutants with concomitant loss of kidney progenitors, and the number of *spi1*-positive cells are increased, consistent with skewing toward primitive myelopoiesis. In contrast, *stag2b* mutants have reduced haematopoietic/vascular mesoderm and downregulation of primitive erythropoiesis. Our results suggest that Stag1 and Stag2 proteins cooperate to balance the production of primitive haematopoietic/vascular progenitors from mesoderm.

## Introduction

Cohesin is a large multi-subunit protein complex that was originally characterised for its role in sister chromatid cohesion during mitosis (Losada, 2008; Onn et al., 2008; Nasmyth and Haering, 2009). Cohesin subunits Smc1A, Smc3 and Rad21 form a large ring-shaped structure that entraps and holds together DNA strands (Nasmyth, 2011). A fourth subunit of either Stag1 or Stag2 binds to cohesin by contacting Rad21 and Smc subunits (Shi et al., 2020), and is required for the association of cohesin with DNA.

Additional roles for cohesin include DNA damage repair and the control of gene expression (Dorsett and Strom, 2012). The gene expression function of cohesin is thought to derive from cohesin’s role in three-dimensional genome organisation (Bonora et al., 2014; Rowley and Corces, 2018). Together with the zinc finger protein, CCCTC-binding factor (CTCF), cohesin organizes the genome into large loops known as topologically-associating domains (TADs) (Vietri Rudan and Hadjur, 2015; Hnisz et al., 2016; Rowley and Corces, 2018). The current theory is that cohesin forms loops by extrusion of DNA through the cohesin ring, and CTCF bound in convergent orientation limits extrusion to delineate loop size (Fudenberg et al., 2018; Hansen, 2020).

Inside TADs, cohesin can mediate smaller loops that link genes to their regulatory elements (Merkenschlager and Odom, 2013). Differential formation of sub-TAD gene regulatory loops is thought to be key to cell type specification during development (Hnisz et al., 2016). Several previous studies have linked mutations in cohesin subunits with tissue-specific changes in gene expression (Dorsett, 2009; Merkenschlager, 2010; Horsfield et al., 2012; Kawauchi et al., 2016). Therefore, via its role in genome organization, cohesin plays a crucial role in developmental gene expression.

Germline mutations in genes encoding the cohesin loader NIPBL, or in cohesin subunits, cause a spectrum of human developmental disorders, the best known of which is Cornelia de Lange Syndrome (CdLS). These disorders, known as “cohesinopathies”, are characterized by multifactorial developmental anomalies, intellectual disability and growth delay (Liu and Krantz, 2009). On the other hand, somatic mutations in cohesin subunits contribute to the development of several types of cancer, including bladder cancer (15-40%), endometrial cancer (19%), glioblastoma (7%), Ewing’s sarcoma (16-22%) and myeloid leukemias (5-53%) (De Koninck and Losada, 2016; Hill et al., 2016; Waldman, 2020). How pathogenicity arises from cohesin mutation is poorly understood, but for both cohesinopathies and cancers, causality is thought to derive primarily from the gene expression function of cohesin rather than its cell division role (Hill et al., 2016; Waldman, 2020).

Notably, there is a particularly high frequency of cohesin gene mutations in myeloid malignancies (Kon et al., 2013; Yoshida et al., 2013; Leeke et al., 2014; Thol et al., 2014; Thota et al., 2014; Papaemmanuil et al., 2016). The high frequency of cohesin mutations in myeloid cancers likely reflects cohesin’s role in determining haematopoietic lineage identity and controlling the differentiation of haematopoietic stem cells (Mazumdar et al., 2015; Mullenders et al., 2015; Viny et al., 2015; Galeev et al., 2016; Viny et al., 2019).

Several previous studies have investigated the role of cohesin in animal development. In Drosophila, Nipped-B and cohesin control *cut* gene expression in the wing margin (Dorsett et al., 2005) and mutations in *Nipped-B* or cohesin genes have dosage-dependent effects on the expression of developmental genes (Dorsett, 2009; Gause et al., 2010). In mice, deficiency in Nipbl or cohesin subunits results in multifactorial developmental abnormalities that mimic CdLS (Kawauchi et al., 2009; Remeseiro et al., 2012b; Smith et al., 2014; Newkirk et al., 2017). Zebrafish models show that Nipbl and cohesin are important for tissue-specific gene regulation (Monnich et al., 2009; Rhodes et al., 2010; Muto et al., 2011), including expression of *hox* genes (Muto et al., 2014) and *runx* genes (Horsfield et al., 2007).

Although animal models have been crucial to understanding the developmental origins of both cohesinopathies (Kawauchi et al., 2016) and haematological malignancies (Viny and Levine, 2018), much remains to be discovered. It is still unclear how cohesin contributes to cell type specification in early development and cell lineage specification. Furthermore, whether all the protein components of cohesin operate equivalently in development is undetermined.

In zebrafish, a forward genetic screen determined that mutation in cohesin subunit *rad21* led to loss of *runx1* expression in the posterior lateral plate mesoderm of zebrafish embryos during early somitogenesis. Knock down of the Smc3 subunit of cohesin also eliminated mesoderm *runx1* expression (Horsfield et al., 2007). Runx1 is essential for definitive haematopoiesis, and is itself affected by mutations and translocations in myeloid malignancies (Downing et al., 2000; Speck, 2001). Previous research shows that *runx1* is directly regulated by Rad21-cohesin in zebrafish (Horsfield et al., 2007; Marsman et al., 2014) and leukaemia cell lines (Antony et al., 2020). Loss of mesoderm-expressed *runx1* at the very earliest time of blood development in *rad21* mutants suggests that the onset of haematopoietic differentiation from the mesoderm might require functional cohesin.

The Stag subunits differ from core cohesin subunits Rad21, Smc1 and Smc3 in that they have redundant roles in cell division, such that a complete loss of Stag2 is tolerated due to partial compensation by Stag1. In addition, Stag1 preferentially associates with CTCF to organise TADs whereas Stag2 mediates short-range cell-specific interactions (van der Lelij et al., 2017; Liu et al., 2018; Cuadrado and Losada, 2020). In mice, homozygous loss of Stag1 is lethal at embryonic day 11.5 (E11.5) (Remeseiro et al., 2012a). While adult loss of Stag2 is tolerated, homozygous Stag2-null mouse embryos die by mid-gestation with developmental delay and defective heart morphogenesis (De Koninck et al., 2020). When Stag2 is ablated somatically in adults, increased self-renewal of HSCs accompanied by myeloid skewing is observed (Viny et al., 2019; De Koninck et al., 2020). However, early lethality of *Stag* mutations makes investigating the embryonic function of Stag1/Stag2 cohesin difficult in mammalian models.

In this study, we characterised the expression of Stag paralogues in early zebrafish development, and investigated whether, like Rad21, cohesin Stag subunits affect haematopoietic differentiation from mesoderm in zebrafish embryos.

## Results

### Evolution and embryonic expression of zebrafish Stag paralogues

Zebrafish have four gene paralogues encoding Stag proteins: *stag1a*, *stag1b*, *stag2a* and *stag2b.* To determine if these paralogues are likely to be functional, we characterised their evolutionary conservation and expression in zebrafish embryos.

Phylogenetic analysis of Stag protein sequences using the PhyML algorithm segregated Stag1 and Stag2 into distinct clusters. Stag2b clustered more closely with other vertebrate Stags than Stag2a, while the two Stag1 paralogues had similar levels of divergence (Figure 1A). Whole-mount *in situ* hybridisation (WISH) (Figure 1B) and quantitative RT-PCR (qPCR) (Figure 1C,D) was then used to analyse expression of the four *stag* paralogues in zebrafish embryogenesis.

**Figure 1.**
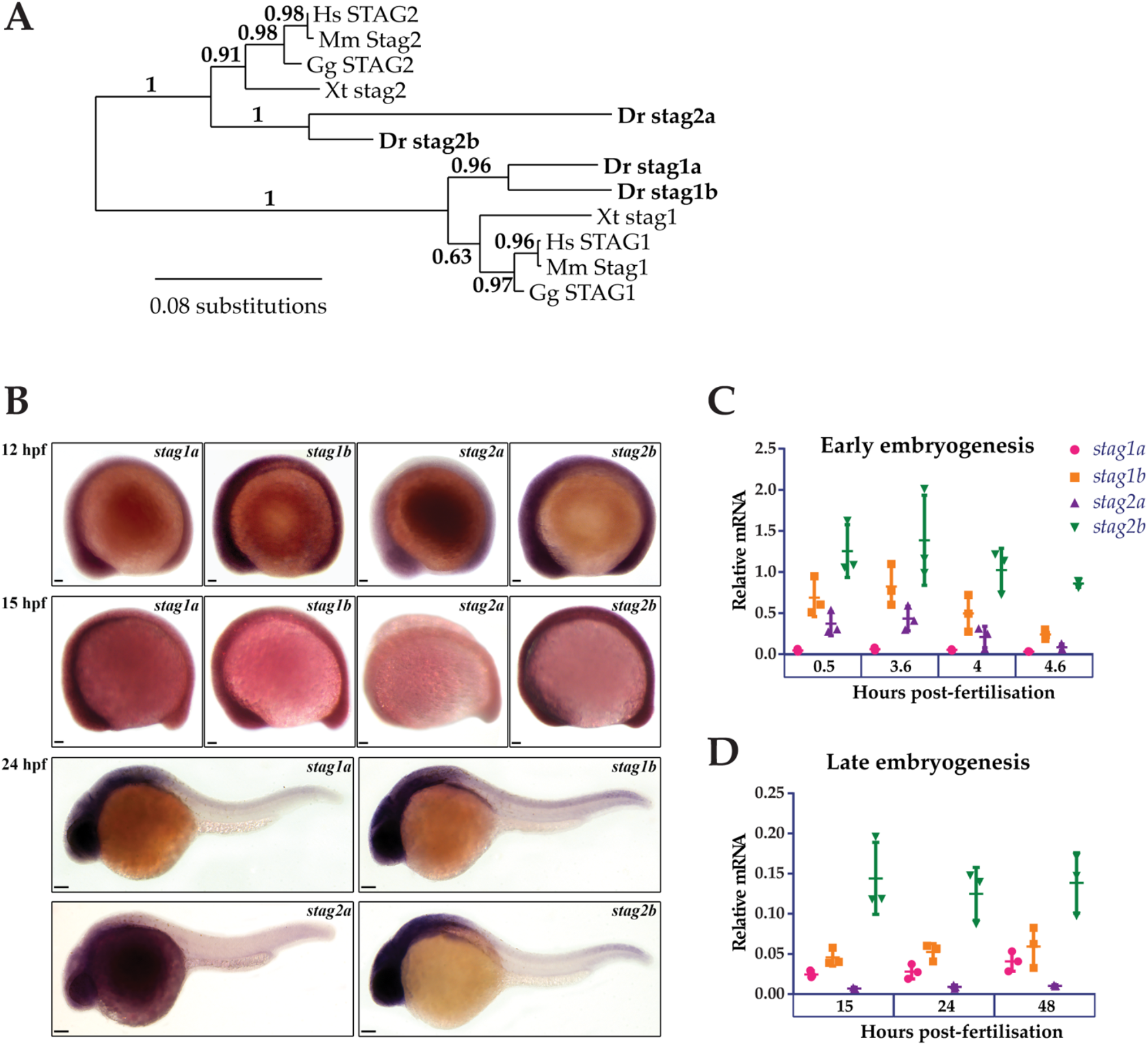
Phylogenetic analysis and embryonic expression of Stag paralogues. **(A)** Phylogenetic analysis of predicted protein sequences using the maximum likelihood approach. The accession numbers for the protein sequences used in this analysis are listed in Supplementary Table 1. **(B)** Whole-mount *in situ* hybridisation of *stag1a, stag1b, stag2a and stag2b* during early embryogenesis. Lateral views are shown, anterior to the left. Scale bars are 50 μm for embryos at 12 hpf and 15 hpf and 100 μm for embryos at 24 hpf. **(C,D)** mRNA expression of *stag* paralogues at the indicated time points during **(C)** early embryogenesis and **(D)** late embryogenesis. Each data point represents mRNA isolated from a pool of 30 embryos. Graphs are mean +/− one standard deviation. Expression was normalised to the reference genes, b-actin and rpl13a (Supplementary Figure 1A)

At early gastrula stages, all four *stag* paralogues showed ubiquitous expression although *stag2a* expression was noticeably reduced compared with *stag1a/b* and *stag2b*. By 24 hours post-fertilisation (hpf), expression of *stag1a/b* and *stag2b* was robust in anterior regions with high cellular density, similar to that observed for genes encoding other cohesin subunits (Monnich et al., 2009), while *stag2a* was barely expressed above background (Figure 1B).

We used qPCR to quantify mRNA expression of the *stag* paralogues at different embryonic timepoints. All four paralogues were both maternally deposited and zygotically expressed with *stag1b* and *stag2b* being the most expressed throughout embryogenesis. Notably, *stag1a* was predominantly zygotically expressed whereas *stag2a* showed maternal deposition that was downregulated post-midblastula transition (Figure 1C,D).

In summary, all four Stag paralogues are expressed during development, indicating that they have potential to be functional.

### Generation of *stag1* and *stag2* mutant zebrafish lines

To determine the physiological roles of the four paralogues, we generated loss-of function germline zebrafish mutants in individual *stag* genes. CRISPR guide RNAs (Supplementary Table 2) were designed to truncate the Stag paralogues upstream of the STAG domain, which spans exons 6 and 7 in all paralogues. We recovered the following germline mutations: 38 bp insertion in exon 3 of *stag1a*, 13 bp deletion in exon 3 of *stag1b* and 7 bp deletion in exon 3 of *stag2b* (Figure 2A and Supplementary Figure 2). No germline mutations could be recovered in *stag2a* despite evaluating multiple guide RNAs. The three zebrafish *stag* mutant alleles we successfully generated were named *stag1a*^*nz204*^, *stag1b*^*nz205*^, and *stag2b*^*nz207*^.

**Figure 2.**
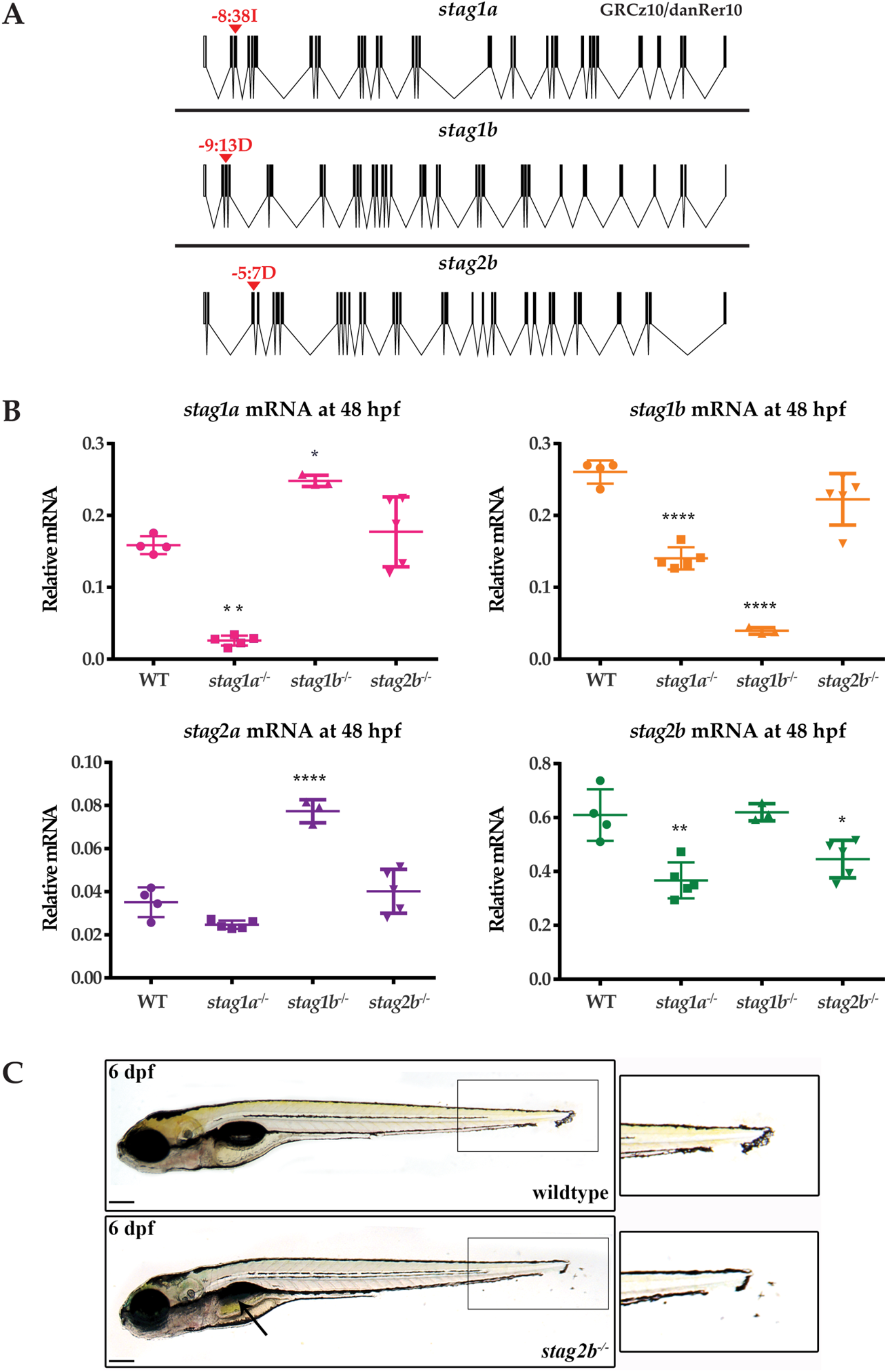
Generation of zebrafish *stag* germline mutants. CRISPR-Cas9 genome editing was used to generate germline mutations in *stag1a*, *stag1b* and *stag2b*. **(A)** Exon diagrams of the respective paralogues showing details of the editing strategy. sgRNA binding sites are marked by red arrowheads with the type of mutation generated indicated above. **(B)** mRNA levels of the four paralogues in each of the mutant lines indicated on the x-axis. Each data point represents mRNA isolated from a pool of 30 embryos. All graphs are mean +/− one standard deviation. * P ≤ 0.05, ** P ≤ 0.01, *** P ≤ 0.001, **** P ≤ 0.0001; one-way ANOVA. Expression was normalised to the reference gene, *b-actin* (Supplementary Figure 1C) **(C)** *stag2b^nz207^* mutants have displaced pigment cells in the tail fin, zoom-ins are shown in insets. Mutants also show mild developmental delay with late swim bladder inflation as indicated by the black arrow. Scale bars are 200 μm.

To confirm knockdown and to evaluate paralogue compensation, we measured the mRNA levels of the four paralogues at 48 hpf using qPCR (Figure 2B). In *stag1a*^*nz204*^ mutants, *stag1a* mRNA was significantly reduced and was accompanied by significant downregulation of *stag1b* and *stag2b* mRNA. In *stag1b*^*nz205*^ mutants, *stag1b* mRNA was significantly reduced and was accompanied by a significant upregulation of *stag1a* and *stag2a* mRNA levels, indicating potential transcriptional compensation. In *stag2b*^*nz207*^ mutants, *stag2b* mRNA was modestly but significantly reduced with no changes in the other paralogues.

The *stag1a^nz204^*, *stag1b^nz205^*, and *stag2b*^*nz207*^ zebrafish mutants were all homozygous viable to adulthood, and fertile. While *stag1a*^*nz204*^ mutants had no apparent larval phenotype, both *stag1b*^*nz205*^ and *stag2b*^*nz207*^ mutants exhibited mild developmental delay. In addition, *stag2b*^*nz207*^ mutants had displaced pigment cells in the tail fin by 54 hpf with a penetrance of ~80-85% (Figure 2C). Injection of 200 pg functional *stag2b* mRNA in *stag2b*^*nz207*^ mutants rescued the displaced pigment cells (Supplementary Figure 3A).

Despite the presence of a 7 bp deletion in the *stag2b* gene in *stag2b*^*nz207*^ mutants, downregulation of the *stag2b* transcript was rather modest (Figure 2B). Therefore, we sought to confirm loss of function in *stag2b*^*nz207*^ mutants by determining if a morpholino oligonucleotide targeting *stag2b* phenocopies the *stag2b*^*nz207*^ mutation. Injection of 0.5 mM *stag2b* morpholino generated the same pigment cell displacement phenotype that was observed in the *stag2b*^*nz207*^ mutant with no observable toxicity. Furthermore, injection of 0.5 mM *stag2b* morpholino into *stag2b*^*nz207*^ embryos caused no additional abnormalities (Supplementary Figure 3B). These observations indicate that the *stag2b*^*nz207*^ allele is likely to be a true loss of function.

Overall, it appears that three of the Stag paralogues, Stag1a, Stag1b and Stag2b, are individually dispensable for zebrafish development and reproduction. We were not able to recover zebrafish mutant for *stag2a*; its early maternal expression supports the idea that this subunit may be essential in the germline.

### *Stag* mutations reduce primitive erythroid cells in 24 hpf zebrafish embryos

Previously, we found that a nonsense mutation in cohesin subunit *rad21* inhibits primitive erythropoiesis and blocks the emergence of differentiated myeloid cells (Horsfield et al., 2007). Therefore, we were interested to determine if cohesin *stag* subunit mutations also affect haematopoiesis in zebrafish embryos. Whole mount *in situ* hybridisation (WISH) was used to determine if expression of markers of primitive and definitive haematopoiesis is affected in zebrafish *stag* mutants at 24 hpf and 36 hpf. We found that *stag1b*^*nz205*^ mutants had no haematopoietic phenotype (data not shown), but that the *stag1a*^*nz204*^ and *stag2b*^*nz207*^ mutations both had modest effects on embryonic haematopoiesis.

Expression of *gata1* marks primitive erythroid cells, and expression of *spi1* (also known as *pu.1*), primitive myelopoiesis. Expression of *gata1* at 24 hpf was downregulated in *stag1a*^*nz204*^ and *stag2b*^*nz207*^ homozygous and heterozygous mutants, indicating loss of primitive erythroid cells. *gata1* expression was rescued by injection of functional *stag1a* or *stag2b* mRNA (Figure 3A). In contrast, we found that *stag1a* and *stag2b* mutation had divergent effects on primitive myelopoiesis: *stag1a*^*nz204*^ increased *spi1* expression, while *stag2b*^*nz207*^ had no effect (Figure 3B). The results suggest that Stag1a and Stag2b promote *gata1*-mediated primitive erythropoiesis and in addition, Stag1a restricts *spi1*-mediated primitive myelopoiesis.

**Figure 3.**
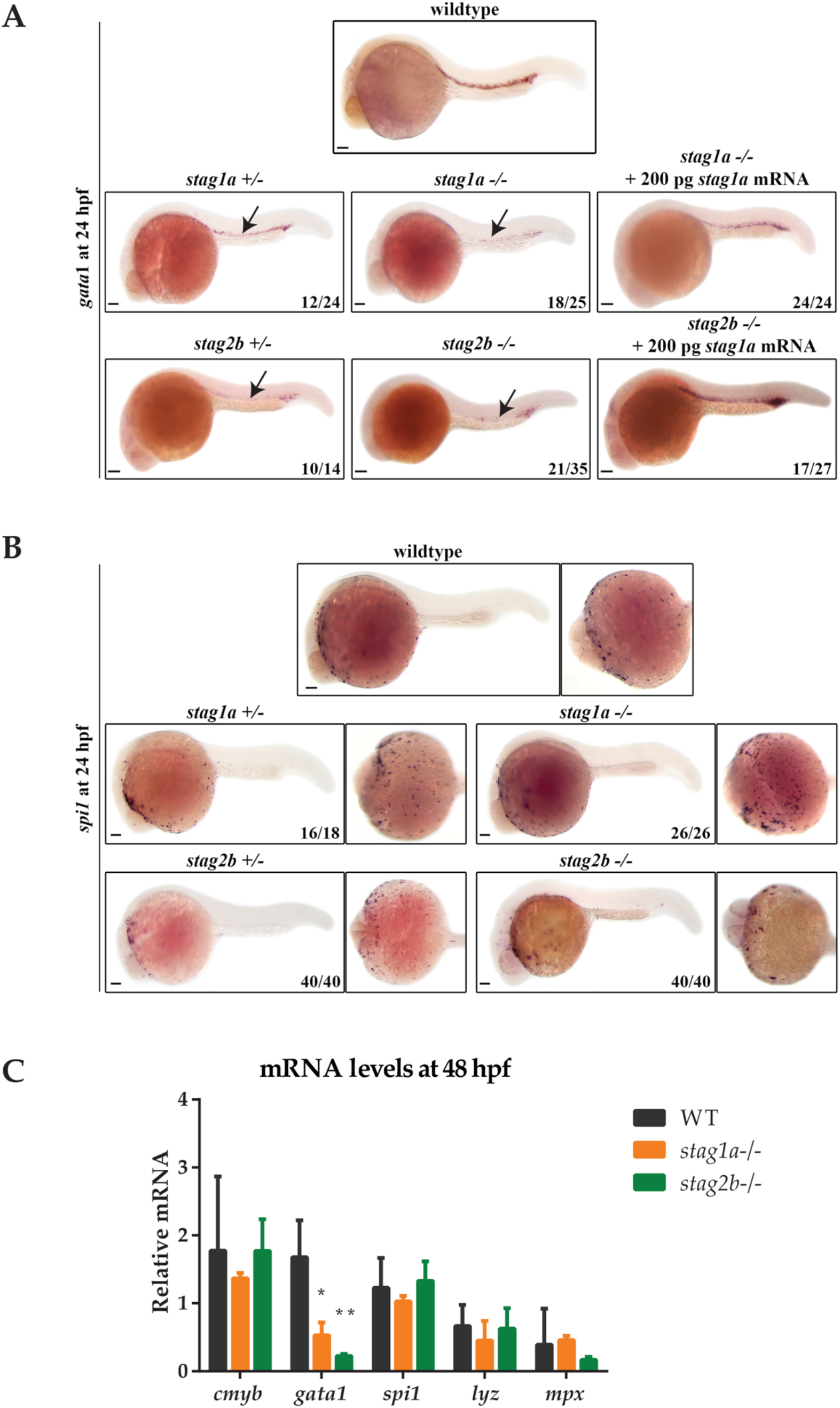
*Stag* mutations alter the number of *gata1-* and *spi1*-positive cells in 24 hpf zebrafish embryos. **(A)** Lateral views of *gata1* expression in whole-mount embryos at 24 hpf; anterior to the left. *gata1* expression is reduced in *stag1a*^*nz204*^+/− and *stag1a*^*nz204*^−/− embryos and is rescued upon injection of functional *stag1a* mRNA. *gata1* expression is reduced in *stag2b*^*nz207*^+/− and *stag2b*^*nz207*^−/− embryos and is rescued upon injection of functional *stag2b* mRNA. Reduced expression is indicated by arrows. **(B)***spi1* expression in whole-mount embryos at 24 hpf. Left panels show lateral views and right panels show ventral views; anterior to the left. The number of *spi1*-positive cells is increased in *stag1a*^*nz204*^+/− and *stag1a*^*nz204*^−/− embryos. *spi1* expression in *stag2b*^*nz207*^ heterozygous embryos and *stag2b*^*nz207*^ homozygous mutant embryos is comparable to wildtype. Scale bars are 100 μm. The number of embryos is indicated in lower-right-hand corners. **(C)** mRNA levels of haematopoietic stem cell marker *cmyb* and erythroid or myeloid lineage markers in *stag1a*^*nz204*^ and *stag2b*^*nz207*^ homozygous mutant embryos at 48 hpf. The bar graph shows the mean +/− one standard deviation. * P ≤ 0.05, ** P ≤ 0.01; one-way ANOVA. Expression was normalised to the reference gene, *b-actin*.

Definitive haematopoietic stem cells (HSCs) in the ventral wall of the dorsal aorta are marked by *runx1* and *cmyb* expression at 36 hpf. HSC expression of *runx1* was moderately reduced in *stag1a*^*nz204*^ mutants and unchanged in *stag2b*^*nz207*^ mutants (Supplementary Figure 4). Quantitative PCR of RNA isolated from 48 hpf *stag1a*^*nz204*^ and *stag2b*^*nz207*^ embryos showed that transcript levels of *cmyb, mpx* and *lyz* mRNA were similar between mutants and wild type, indicating that definitive myelopoiesis is intact in the mutants. In contrast, *gata1* expression remained reduced in both *stag1a*^*nz204*^ and *stag2b*^*nz207*^ mutants at 48 hpf (Figure 3C), indicating that the deficiency in erythropoiesis is sustained from early development. Therefore, Stag1a and Stag2b appear to promote erythropoiesis during embryonic haematopoiesis, but are dispensable for myelopoiesis.

### Stag1a and Stag2b are important for specification of *scl*-positive cells in the haematopoietic mesoderm

A null cohesin *rad21* mutation causes a striking, complete loss of *runx1* expression in the posterior lateral mesoderm (PLM) of zebrafish embryos at 4-15 somite stages (Horsfield et al., 2007). This observation prompted us to investigate whether *stag* mutations also affect expression of *runx1* and other lineage-defining genes in the intermediate mesoderm.

WISH with a riboprobe detecting *runx1* expression in the PLM on 15 hpf embryos (14 somites) revealed that *stag1a*^*nz204*^ and *stag2b*^*nz207*^ mutants had relatively normal PLM *runx1* expression (Figure 4A). We observed minor expansion in the PLM domain of *runx1* in *stag1a*^*nz204*^ mutants, and minor localised reduction of *runx1* expression in *stag2b*^*nz207*^ mutants; however, qPCR revealed that total *runx1* transcript levels are not significantly different between mutants and wild type (Figure 4C). Therefore, unlike *rad21* mutation, *stag1a* or *stag2b* mutations are by themselves not sufficient to cause dramatic changes to *runx1* expression.

**Figure 4.**
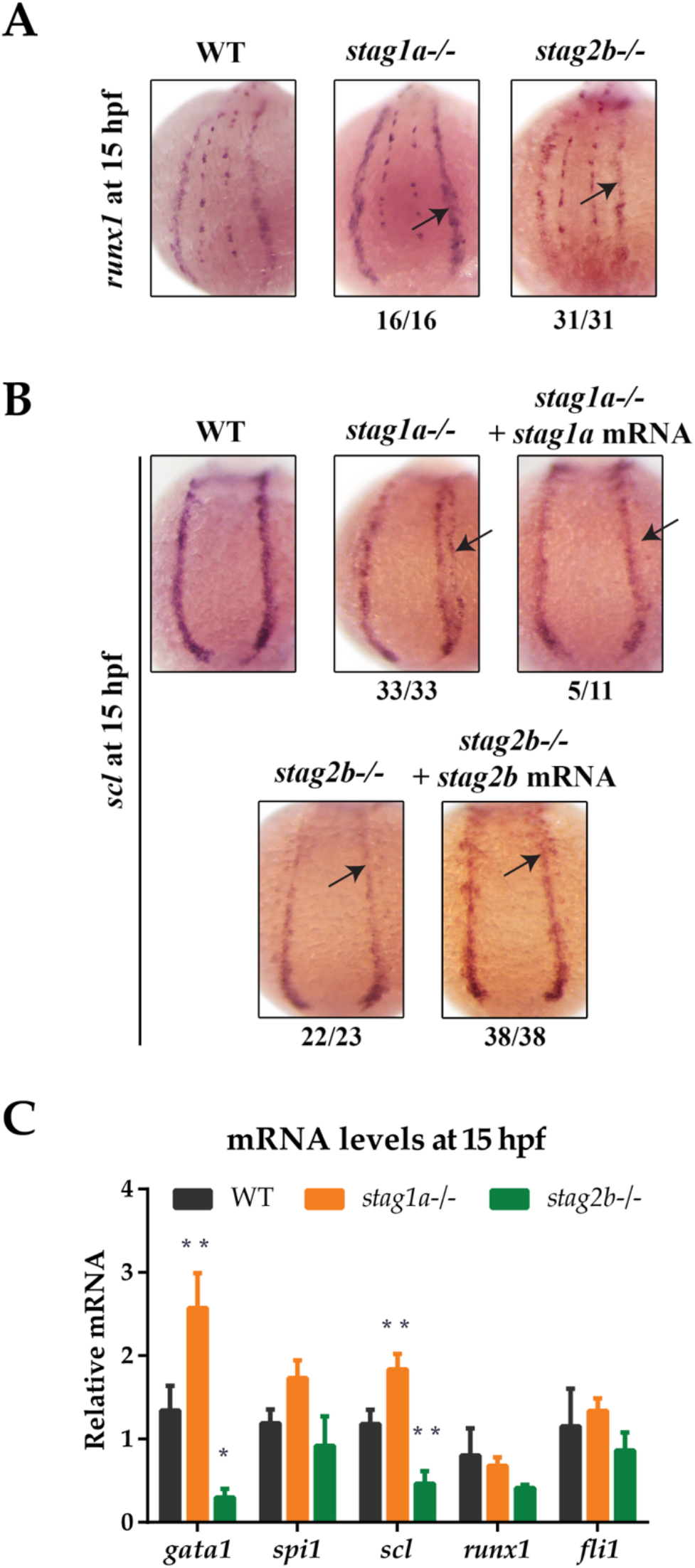
*stag1a* and *stag2b* mutations alter the number of *scl*-positive cells in the posterior lateral mesoderm at 15 hpf. **(A)** *runx1* expression in wholemount embryos at 15 hpf. Posterior views of the PLM are shown; dorsal to the top. In *stag1a*^*nz204*^ homozygous mutant embryos, runx1 expression is slightly increased. In *stag2b*^*nz207*^ homozygous mutant embryos, *runx1* expression is slightly reduced. Changes in expression are marked by arrows and the number of embryos is indicated below each panel. **(B)** *scl* expression in whole-mount embryos at 15 hpf. Posterior views of the PLM are shown; dorsal to the top. In *stag1a*^*nz204*^ homozygous mutant embryos, expanded expression of *scl* laterally into the PLM is dampened upon injection of functional *stag1a* mRNA. In *stag2b*^*nz207*^ homozygous mutant embryos, *scl* expression is reduced in the anterior PLM and is rescued upon injection of functional *stag2b* mRNA. Changes in expression are marked by arrows and the number of embryos is indicated below each panel. **(C)** mRNA levels of mesoderm-derived haematopoietic and endothelial markers at 15 hpf. The bar graph shows the mean +/− one standard deviation. * P ≤ 0.05, ** P ≤ 0.01; oneway ANOVA. Expression was normalised to the reference genes, *b-actin* and *rpl13a* (Supplementary Figure 1B).

Expression of the *scl* (*tal-1*) gene marks a subset of cells in the PLM that will later go on to assume either vascular or haematopoietic identity. Surprisingly, we observed significant differences in the expression pattern of *scl* in the PLM of *stag1a*^*nz204*^ and *stag2b*^*nz207*^ mutants at 15 hpf (14 somites) (Figure 4B). An expanded lateral domain of *scl* expression appeared in the PLM of *stag1a*^*nz204*^ mutants, and was rescued by injection of *stag1a* mRNA (Figure 4B). In contrast, *scl* expression was reduced in the anterior PLM of *stag2b*^*nz207*^ mutants, and this was rescued by injection of *stag2b* mRNA (Figure 4B). The observed changes in *scl* expression were reinforced by qPCR analysis (Figure 4C), which showed an increase of *scl* transcript in *stag1a*^*nz204*^ and decrease in *stag2b*^*nz207*^ mutants, respectively. In contrast to observations in 24 hpf embryos, *gata1* transcript levels were increased in *stag1a*^*nz204*^ mutants along with a slight increase in *spi1* mRNA. Expression of the vascular marker, *fli1*, was not significantly altered (Figure 4C).

The results suggest that during early somitogenesis in *stag1a*^*nz204*^ mutants, *scl*-positive cell numbers are expanded and accompanied by the upregulation of primitive haematopoietic markers. In contrast, both *scl* and *gata1* are downregulated in *stag2b*^*nz207*^ mutants suggesting a reduction in *scl*-positive haematopoietic/vascular progenitors.

### Loss of Stag1a, but not Stag2b, alters gene expression domains in the posterior lateral mesoderm

During early somitogenesis, the PLM contains non-overlapping stripes of *pax2a*-expressing pronephric progenitors adjacent to the *scl*-expressing cells. We were curious to know whether changes in the *scl*-positive population in the *stag* mutants influenced adjacent cell populations, such as pronephric progenitors, in the mesoderm.

At 12 hpf (10 somites), *scl* expression was expanded in *stag1a*^*nz204*^ mutants, while in *stag2b*^*nz207*^ mutants, *scl* expression was slightly reduced (Figure 5A). This is finding is consistent with observations of 15 hpf embryos (Figure 4B,C; Supplementary Figure 5A). Notably, the PLM zone of *pax2a* expression was reduced concomitant with expansion of *scl*-expressing cells in the PLM of *stag1a*^*nz204*^ mutants (Figure 5B; Supplementary Figure 5B). These results suggest that *scl*-positive haematopoietic/endothelial progenitors are expanded at the expense of pronephric progenitors in *stag1a*^*nz204*^ mutants. In contrast, in *stag2b*^*nz207*^ mutants with reduced *scl* transcript, expression of *pax2a* was maintained in the PLM but reduced in the optic stalk compared with wild type (Figure 5B,C; Supplementary Figure 5B).

**Figure 5.**
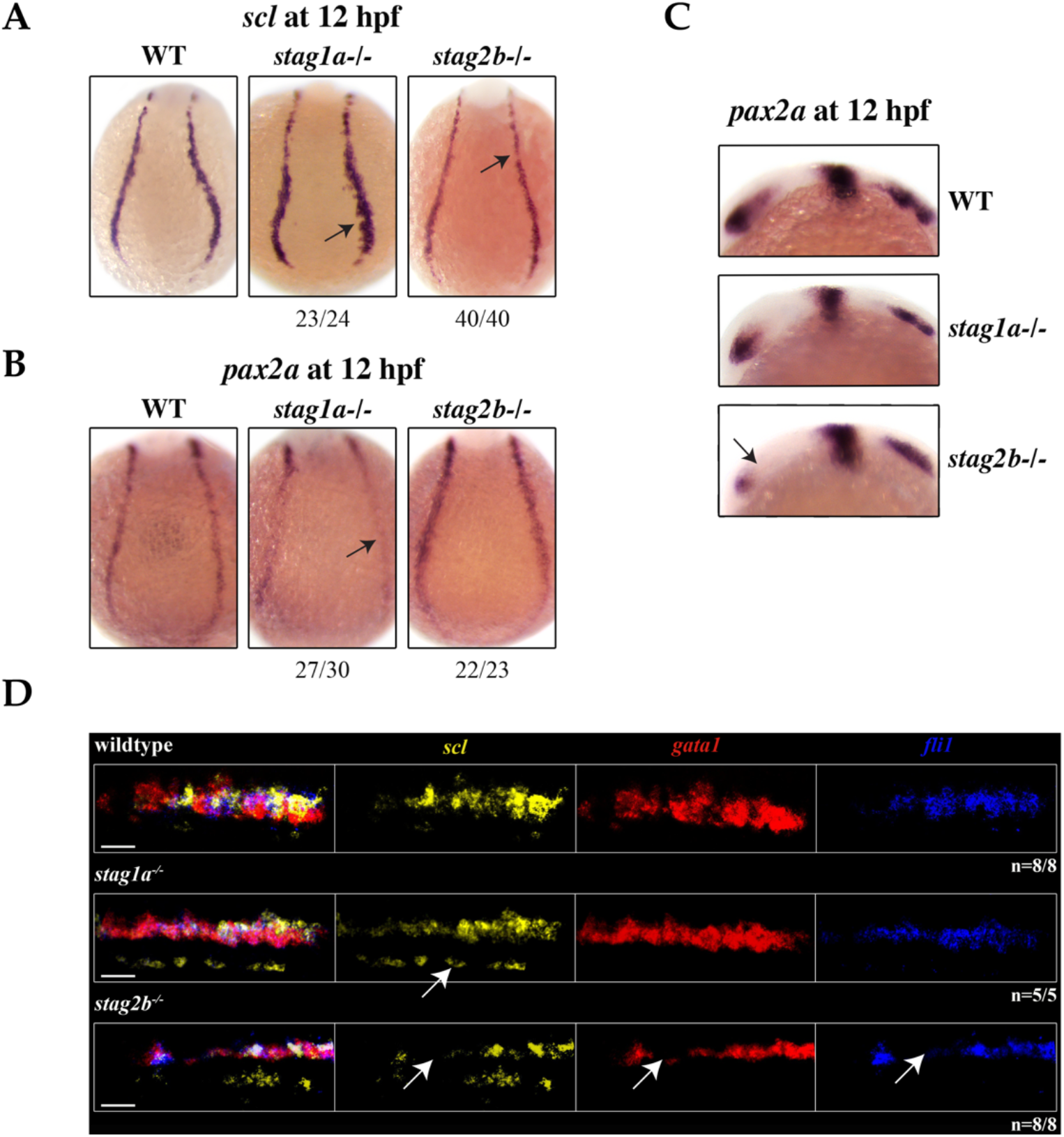
*stag1a* and *stag2b* mutations affect cell identity in the posterior lateral mesoderm at 12 hpf. **(A)** *scl* expression in whole-mount embryos at 12 hpf, posterior views of the PLM; dorsal to the top. In *stag1a*^*nz204*^ homozygous mutant embryos *scl* expression is expanded. In *stag2b*^*nz207*^ homozygous mutant embryos, *scl* expression is reduced. Changes in expression are marked by arrows and the number of embryos is indicated below each panel. **(B)** *pax2* expression in whole-mount embryos at 12 hpf, posterior views of the PLM; dorsal to the top. In *stag1a*^*nz204*^ homozygous mutant embryos, *pax2* expression in the PLM is markedly reduced. In *stag2b*^*nz207*^ homozygous mutant embryos, *pax2* expression is comparable to wild type. Changes in expression are marked by arrows and the number of embryos is indicated below each panel. **(C)***pax2* expression in whole-mount embryos at 12 hpf, lateral views of the head region; anterior to the left. Anterior *pax2* expression is specifically reduced in the optic stalk of *stag2b*^*nz207*^ homozygous mutant embryos. **(D)**Multiplexed *in situ* HCR of *scl* (Alexa Fluor 488, false colour yellow), *gata1* (Alexa Fluor 594, false colour red) and *fli1* (Alexa Fluor 647, false colour blue) expression at 15 hpf. High magnification maximum intensity projections of a single PLM stripe; posterior views with anterior to the left. Expression domains of *scl* broadly overlap *gata1* and *fli1* in all embryos. Ectopic *scl* expression, indicated by white arrow, in *stag1a*^*nz204*^ homozygous mutant embryos does not overlap *gata1* or *fli1* expression domains. In *stag2b*^*nz207*^ homozygous mutant embryos, expression of all three markers is reduced. Scale bars are 10 μm. The number of embryos analysed is indicated below the respective panels.

A subset of *scl*-positive cells also express *gata1* and acquire a haematopoietic fate while the remaining cells express *fli1* acquiring an endothelial fate. We next wanted to determine whether *scl*-positive cells are skewed towards a haematopoietic or vascular fate in the *stag* mutants. Multiplex *in situ* hybridisation using HCR revealed that the expression of *gata1* and *fli1* largely overlap that of *scl* in the PLM (Figure 5D; Supplementary Figure 5A). Ectopic *scl* expression seen in *stag1a*^*nz204*^ mutants did not overlap *gata1* or *fli1* expression, but *gata1* expression appeared more intense than wild type, consistent with qPCR results (Figure 5D; Figure 4C; Supplementary Figure 5A). We detected no differences in the relative composition of *scl*^+^/*gata1*^+^ and *scl*^+^/*fli1*^+^ cells in the PLM of mutants (Supplementary Figure 5A). The results suggest that in *stag1a*^*nz204*^ mutants, expanded *scl* expression does not appear to skew cell fate in the PLM, but transiently increases *gata1* expression.

In *stag2b*^*nz207*^ mutants, the expression domains of *scl*, *gata1* and *fli1* was reduced in the PLM (Figure 5D; Supplementary Figure 5A). Cell composition of the PLM was unchanged in *stag2b*^*nz207*^ mutants (Supplementary Figure 5A), suggesting that reduced *scl*, *gata1* and *fli1* does not influence PLM cell fate.

### Stag1a or Stag2b loss differentially affects the production of primitive myeloid cells in the anterior lateral mesoderm

We next asked if *stag* mutants also affect haematopoietic cell specification in the anterior lateral mesoderm (ALM), a site of primitive myelopoiesis (Berman et al., 2005). At 12 hpf, *scl* expression in the rostral blood island marks a population of cells fated to become *spi1*-positive myeloid cells or *fli1*-positive endothelial cells.

*scl* expression was normal in the ALM of *stag1a*^*nz204*^ mutants at 12 hpf (Figure 6A) but by 15 hpf *scl* expression was markedly increased in *stag1a*^*nz204*^ (Figure 6B, Supplementary Figure 5C). Increased *scl* expression in the ALM of *stag1a*^*nz204*^ mutants was reversed by injection of functional *stag1a* mRNA, which reduced *scl* expression to below normal. In *stag2b*^*nz207*^ mutants, *scl* expression was reduced in the ALM at both 12 and 15 hpf, and was robustly rescued upon injection of *stag2b* mRNA (Figure 5B).

**Figure 6.**
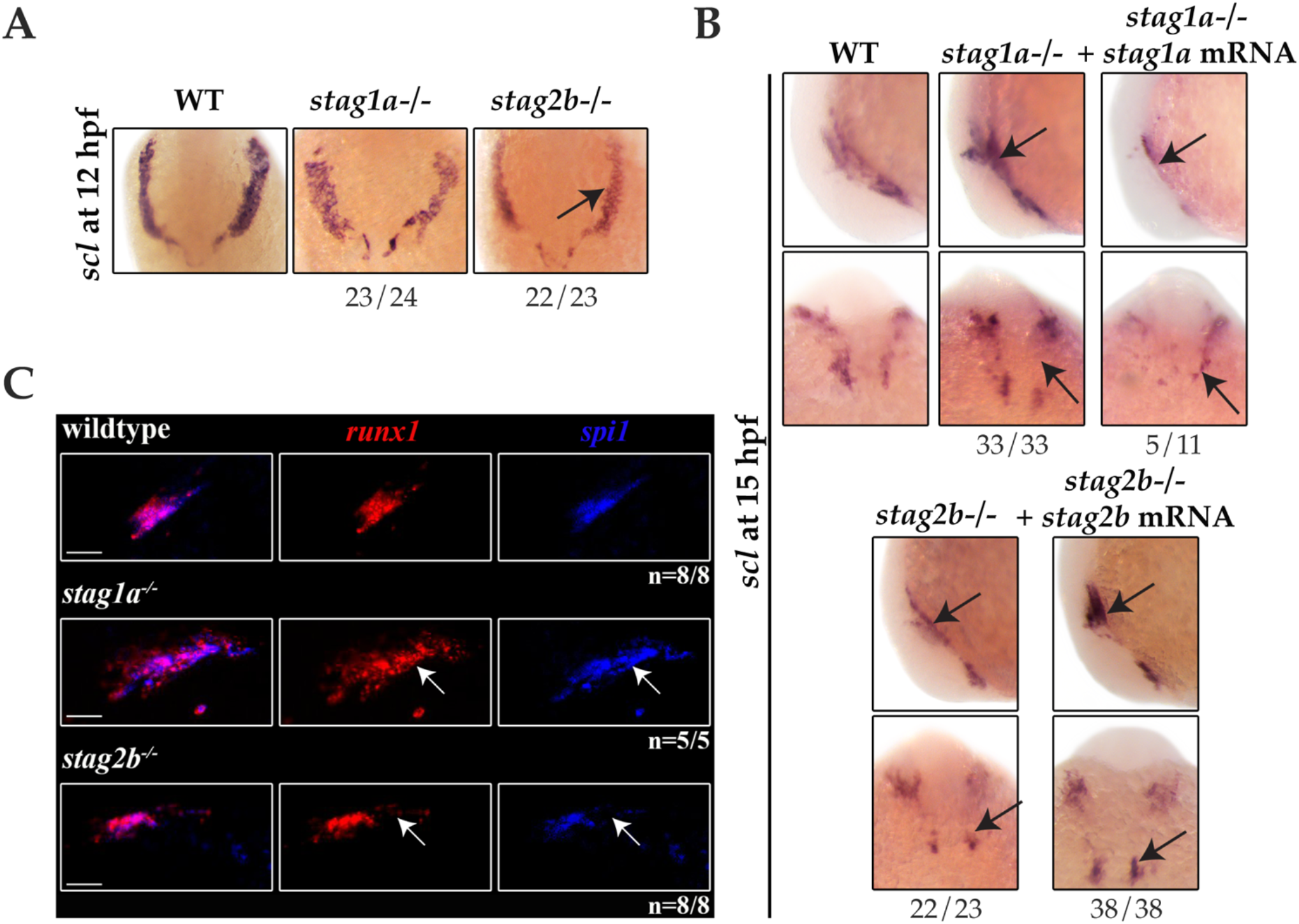
*stag1a* and *stag2b* mutations differentially alter the production of primitive myeloid cells in the anterior lateral mesoderm at 12 hpf. **(A)** *scl* expression in whole-mount embryos at 12 hpf. Ventral views of ALM are shown; dorsal to the top. *scl* expression is comparable to wildtype in *stag1a*^*nz204*^ homozygous mutant embryos and reduced in *stag2b*^*nz207*^ homozygous mutant embryos. **(B)** *scl* expression in whole-mount embryos at 15 hpf. Top panels show lateral views and bottom panels show ventral views of the ALM. Expanded *scl* expression in the ALM of *stag1a*^*nz204*^ mutants is rescued upon injection of functional *stag1a* mRNA. The reduced *scl* expression in the ALM of *stag2b*^*nz207*^ mutants is rescued upon injection of functional *stag2b* mRNA. Changes in expression are marked by arrows and the number of embryos is indicated below each panel. **(C)** Multiplexed *in situ* HCR of *runx1* and *spi1* at 15 hpf. Maximum intensity projections of a single ALM stripe; lateral views with dorsal to the top. Expression domains of *runx1* (Alexa Fluor 647) and *spi1* (Alexa Fluor 514, false colour blue) broadly overlap in all embryos. Both *runx1* and *spi1* are expanded in *stag1a*^*nz204*^ embryos but reduced in *stag2b*^*nz207*^ embryos. Changes in expression are indicated by white arrows. Scale bars are 10 μm. The number of embryos analysed is indicated below the ssrespective panels.

Multiplex HCR expression analysis showed that the population of ALM cells that co-express *runx1* and *spi1* are expanded in the ALM of *stag1a*^*nz204*^ mutants (Figure 6C). In contrast, the same *spi1*/*runx1*-positive ALM population was reduced in *stag2b*^*nz207*^ mutants. Since there was also a modest increase in *spi1*-positive cells in *stag1a*^*nz204*^ mutants at 24 hpf (Figure 3B), these results are consistent with the idea that excess *scl* in *stag1a*^*nz204*^ mutants promotes myelopoiesis in the anterior blood island.

Taken together, the results suggest that in early somitogenesis, Stag1a normally restricts *scl* expression in the ALM and PLM, such that its loss in *stag1a*^*nz204*^ mutants results in a modest expansion of primitive erythroid and myeloid cells at the expense of pronephros specification. In contrast, Stag2b positively regulates the number of *scl*-expressing cells and its loss in *stag2b*^*nz207*^ mutants leads to a reduction of *scl*-derived lineages. However, by 24 hpf *gata1*-positive cells are reduced in both *stag1a*^*nz204*^ and *stag2b*^*nz207*^ mutants, suggesting that erythropenia is a common consequence of an imbalance in *scl*-positive cells. Because both *stag1a*^*nz204*^ and *stag2b*^*nz207*^ mutants are homozygous viable, there must be sufficient redundancies and plasticity to overcome these *stag* mutations in later development.

## Discussion

All four Stag paralogues are expressed in early embryogenesis, suggesting that they are likely to have a function in early development. The maternally and zygotically expressed *stag1b* and *stag2b* are the most abundant of the zebrafish Stags. While zebrafish Stag1a and Stag1b are more or less equally related to mammalian Stag1, the higher zygotic expression of *stag1b* suggests that it is the most predominant isoform in zebrafish. Of the two Stag2 isoforms, *stag2b* is the most abundantly expressed and is also most closely related to mammalian Stag2, suggesting that Stag2b is likely to be the predominant Stag2 in zebrafish.

The *stag2a* paralogue mRNA is present in early embryos up until the mid-blastula transition and then is rapidly downregulated. Interestingly, we detected robust *stag2a* expression in the ovaries of adult zebrafish (Supplementary Figure 6), and little expression elsewhere in adults. It is possible that *stag2a* is required in oocytes for development pre-zygotic genome activation, but is dispensable at later stages. Significantly, we were not able to isolate a CRISPR mutant for *stag2a*, raising the possibility that Stag2a is essential in oocytes its loss does not allow for transmission of a mutation.

All three germline mutations successfully isolated for the Stag paralogues are homozygous viable and fertile, indicating that there is likely to be functional redundancy among Stag proteins throughout development and reproduction. Compensation could be partly transcription based, for example, *stag1b*^*nz205*^ mutant embryos upregulated expression of *stag1a* and *stag2a*. Fish that were mutant for the most abundant Stags, *stag1b*^*nz205*^ and *stag2b^nz207^*, exhibited a slight developmental delay as larvae, and had displaced pigment cells in the tail fin. However, only the *stag1a*^*nz204*^ (which had no morphological phenotype) and *stag2b*^*nz207*^ mutants produced haematopoietic phenotypes in embryos younger than 48 hpf. The sharp increase of *stag1a* expression and the abrupt downregulation of *stag2a* at the mid-blastula transition (leaving *stag2b* as virtually the sole zygotic Stag2) might explain why these two particular mutations caused phenotypes in embryos.

Analysis of primitive haematopoiesis from 24-48 hpf showed that both the *stag1a*^*nz204*^ and *stag2b*^*nz207*^ mutants had a profound decrease in erythroid cells. These findings are in partial agreement with data from mice. Somatic removal of *Stag2* in mice resulted in increased myeloid progenitors and decreased megakaryocyte-erythrocyte progenitors, with consequential myeloid skewing (Viny et al., 2019; De Koninck et al., 2020). However, there is no haematopoietic phenotype in *Stag1*-mutant mice (Viny et al., 2019), contrasting with the erythropenia we observed in zebrafish *stag1a*^*nz204*^ mutants at 24 and 48 hpf.

Although *stag1a*^*nz204*^ and *stag2b*^*nz207*^ mutants both had erythroid deficiency, unexpectedly, only the *stag1a*^*nz204*^ mutant presented with additional early haematopoietic alterations. These included a reduction in *runx1*-positive definitive HSCs at 36 hpf in *stag1a*^*nz204*^ mutants, and striking changes to expression of *scl* in the PLM at 12 and 15 hpf.

The basic helix-loop-helix protein Scl/Tal-1 is expressed in mesoderm and marks both vascular and haematopoietic lineages. Scl is thought to program ventral mesoderm to a haematopoietic fate (Orkin, 1995; Davidson and Zon, 2000; Prummel et al., 2020). Overexpression of zebrafish *scl* leads to an overproduction of blood from mesoderm at the expense of other non-axial mesoderm fates (Gering et al., 1998). Consistent with this, we observed a reduction in expression of *pax2a* in the pronephric mesoderm in *stag1a*^*nz204*^ mutants that had expanded expression of *scl*. However, a concomitant increase in expression of downstream haematopoietic markers *gata1* and *spi1* was only transitory in *stag1a*^*nz204*^ mutants. Expression of *gata1* and *spi1* is increased in 15 hpf *stag1a*^*nz204*^ mutants but by 24 hpf, *spi1* expression was normal and *gata1* expression was reduced.

Stag2 depletion in mice induces both an increase in self-renewal and reduced differentiation capacity in HSCs (Viny et al., 2019). Stag2-deficient mice had downregulation of *spi1* target genes that promote myeloid differentiation. ChIP-sequencing experiments in mice showed that recruitment of Spi1 to genomic binding sites is reduced in the absence of Stag2 (Viny et al., 2019). In zebrafish, loss of Stag2b had little effect on *spi1* expression, but did lead to reduced primitive haematopoiesis overall.

The phenotypes of *stag1a*^*nz204*^ and *stag2b*^*nz207*^ mutants have opposite effects on *scl* expression in early somitogenesis (12 and 15 hpf), but a similar reduction in *gata1*-positive cells by 24 hpf. We suggest that loss of Stag2b leading to reduced *scl* expression limits the pool of progenitors that can contribute to primitive haematopoiesis. Conversely, we propose that increased *scl* expression caused by loss of Stag1a increases haematopoietic progenitors that are subsequently exhausted by early differentiation. These scenarios would explain the erythropenia observed in both *stag1a*^*nz204*^ and *stag2b*^*nz207*^ mutants by 24 hpf (Figure 7).

**Figure 7.**
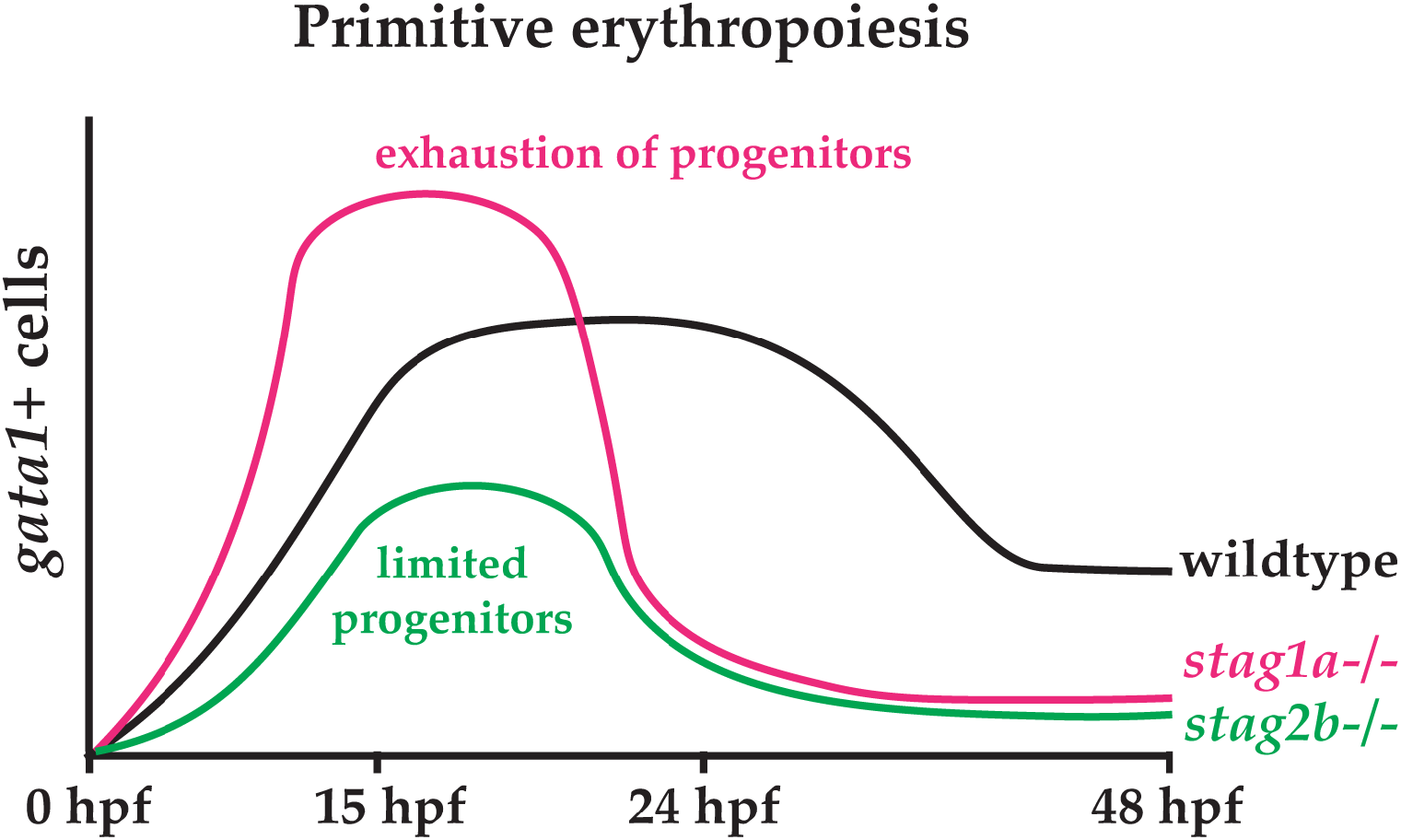
Hypothetical model explaining the effects of Stag1a and Stag2b loss on primitive erythropoiesis. In *stag1a*^*nz204*^ mutants, an expansion of early haematopoietic progenitors driven by increased *scl* expression may lead to precocious differentiation that exhausts the progenitor pool. In *stag2b*^*nz207*^ mutants, a limited pool of haematopoietic progenitors resulting from reduced *scl* expression leads to a reduction in primitive erythropoiesis.

A remaining question is the mechanism by which *stag1a*^*nz204*^ and *stag2b*^*nz207*^ mutants differentially affect *scl* expression. High levels of Bmp signalling induce lateral plate mesoderm and specify haematopoietic fate (Davidson and Zon, 2000; Prummel et al., 2020). Bmp signalling cooperates with Wnt signalling to promote blood fate through activation of homeobox transcription factors Cdx1 and Cdx4 (Lengerke et al., 2008). Previous studies show that mutations in cohesin subunits interfere with canonical Wnt signalling (Avagliano et al., 2017; Chin et al., 2020), so it is possible that loss of Stag1a or Stag2b differentially affect the balance of Bmp and Wnt signalling that directs the production of *scl*-positive cells. Further experimentation will be needed to determine whether this is the case.

In summary, we have characterised the expression and function of zebrafish Stag paralogues in early development and haematopoiesis. We found a surprising role for the Stag1a orthologue in restricting primitive vascular/haematopoietic cell numbers. In contrast, Stag2b loss-of-function reduced progenitor numbers. Subfunctionalisation and homozygous viability of the zebrafish *stag* mutants offer a unique opportunity to dissect cohesin’s developmental functions in the absence of interference from cell cycle phenotypes.

## Materials and Methods

### Zebrafish maintenance

Wild type (WIK) and mutant fish lines were maintained according to established protocols (Westerfield, 1995). Zebrafish procedures were carried out in accordance with the Otago Zebrafish Facility Standard Operating Procedures. Zebrafish mutant lines were developed under AUP-19-17 approved by the University of Otago Animal Ethics Committee. For all experiments, embryos were incubated at 22 °C or 28 °C.

### CRISPR-Cas9 editing

At least three sgRNAs were designed for each *stag* gene using the publicly available CHOPCHOP CRISPR design tool (Montague et al., 2014). sgRNAs were synthesised using a cloning-free approach as previously described (Varshney et al., 2016). Recombinant Cas9 protein was obtained commercially (PNA Bio Inc., Newbury Park, California, USA). Ribonucleoprotein complexes (RNPs) were assembled by mixing sgRNA and Cas9 protein at concentrations of 100 pg/embryo and 300 pg/embryo, respectively in 300 mM KCl. RNPs were incubated for 5 minutes at 37 °C before injection into 1-cell stage WIK embryos. Editing efficiencies were evaluated by genotyping eight embryos from each injection clutch using high resolution melt analysis (HRMA). The most efficient sgRNAs were used to generate germline mutant lines (Supplementary Table 2). Primers used for genotyping are listed in Supplementary Table 3.

### Morpholino and mRNA rescue injections

Morpholinos were purchased from Gene Tools LLC (Philomath, Oregon, USA) for the *stag* genes (Supplementary Table 4). 1-cell stage zebrafish embryos were injected with 0.5 mM of morpholino. Full-length mRNA constructs in pcDNA3.1^+^/C-(K)DYK vectors were obtained from GenScript Biotech (Piscataway, New Jersey, USA) for each *stag* gene. mRNA was synthesised using the mMessage mMachine transcription kit (Ambion, Austin, Texas, USA) and 200 pg was injected into *stag* mutant embryos at the 1-cell stage.

### Whole-mount in situ hybridisation (WISH)

WISH was performed as previously described (Thisse and Thisse, 2008). Digoxigenin-labelled riboprobes for the four *stag* genes were synthesized from PCR clones inserted into pGEM®-T Easy vectors (Promega, Madison, Wisconsin, USA) using T7/Sp6 RNA polymerase (Roche Diagnostics, Basel, Switzerland). Anti-DIG alkaline phosphatase antibody (Roche Diagnostics, Basel, Switzerland) was used for detection, followed by visualization with nitro blue tetrazolium and 5-bromo-4-chloro-3-indolylphosphate (NBT/BCIP) (Roche Diagnostics, Basel, Switzerland). Embryos were imaged using a Leica M205 FA epifluorescence microscope (Leica, Wetzlar, Germany Applications Suite). Primers used for the amplification of *stag* riboprobes are listed in Supplementary Table 3.

### Quantitative PCR (qPCR)

Total mRNA was extracted from pools of 30 embryos using NucleoSpin RNA kit (Macherey-Nagel, Bethlehem, PA, USA). Complementary DNA (cDNA) was synthesized with qScript cDNA SuperMix (Quanta Biosciences, Beverly, MA, USA). Expression levels of the *stag* paralogues (primer sequences in Supplementary Table S3) and haematopoietic markers were measured using SYBR Premix Ex Taq II (Takara Bio Inc., Kusatsu, Japan) on a Roche LightCycler400. Reference genes were *b-actin* and *rpl13a*.

### Hybridisation chain reaction (HCR)

HCR probe sets for *pax2*, *scl*, *runx1*, *gata1*, *spi1* and *fli1* were obtained from Molecular Instruments, Inc. (California, USA). HCR was performed as per the manufacturer’s protocol for zebrafish embryos. Embryos were mounted in 1% agarose and imaged on Nikon C2 confocal microscope (Nikon Corp, Tokyo, Japan NIS-Elements). Image analysis was performed using ImageJ. For embryos shown in figures, maximum intensity projections were generated and brightness/contrast was adjusted with no further processing. For quantitative analysis, individual channels were background-subtracted, auto-thresholded using the RenyiEntropy algorithm (Kapur et al., 1985) and fluorescence intensities were measured. Colocalization analysis was performed using the JACoP plugin (Bolte and Cordelières, 2006) in ImageJ.

### Statistical Analysis

GraphPad PRISM 7 was used for performing all statistical analysis. One-way ANOVAs (Tukey’s multiple comparisons tests) were used for estimating the statistical significance of qPCR and HCR data.

## Author Contributions

SK and JAH designed experiments. SK and AL performed experiments. SK, AL and JAH analyzed data. SK and JAH wrote the paper. All authors read and approved the final manuscript.

## Acknowledgements

The authors would like to thank Noel Jhinku for expert management of the zebrafish facility, and Jisha Antony for helpful advice and discussions.

---

**Supplementary Figure 1.**
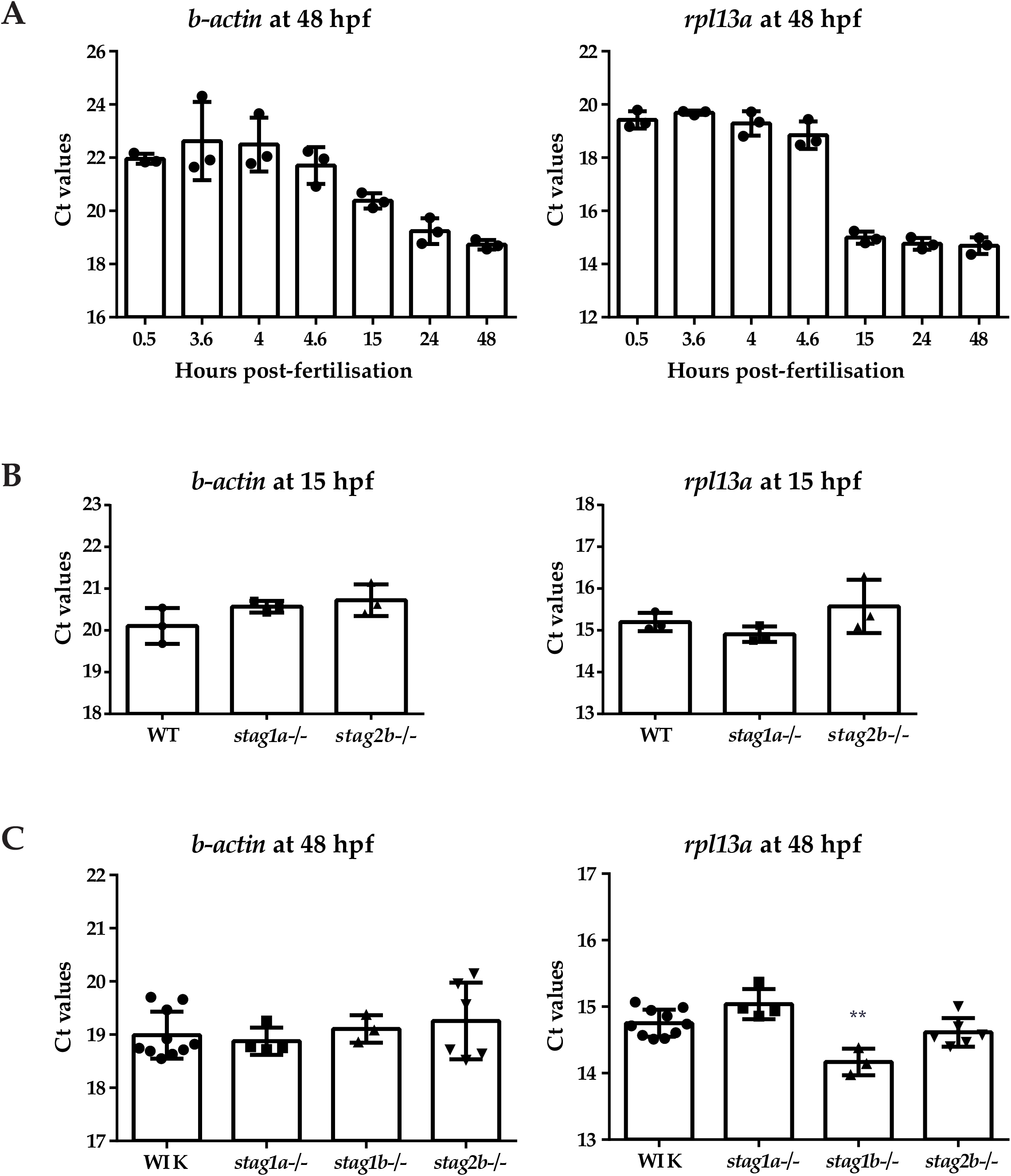
Stability of reference genes used for quantitative RT-PCR (qPCR) normalisation. **(A)** Ct values of *b-actin* and *rpl13a* in wildtype embryos at 48 hpf. **(B)** Ct values of *b-actin* and *rpl13a* in wildtype and mutant embryos at 15 hpf. **(C)** Ct values of *b-actin* and *rpl13a* in wildtype and mutant embryos at 48 hpf. Expression of *rpl13a* is altered in mutants. **P≤0.01; one-way ANOVA.

**Supplementary Figure 2.**
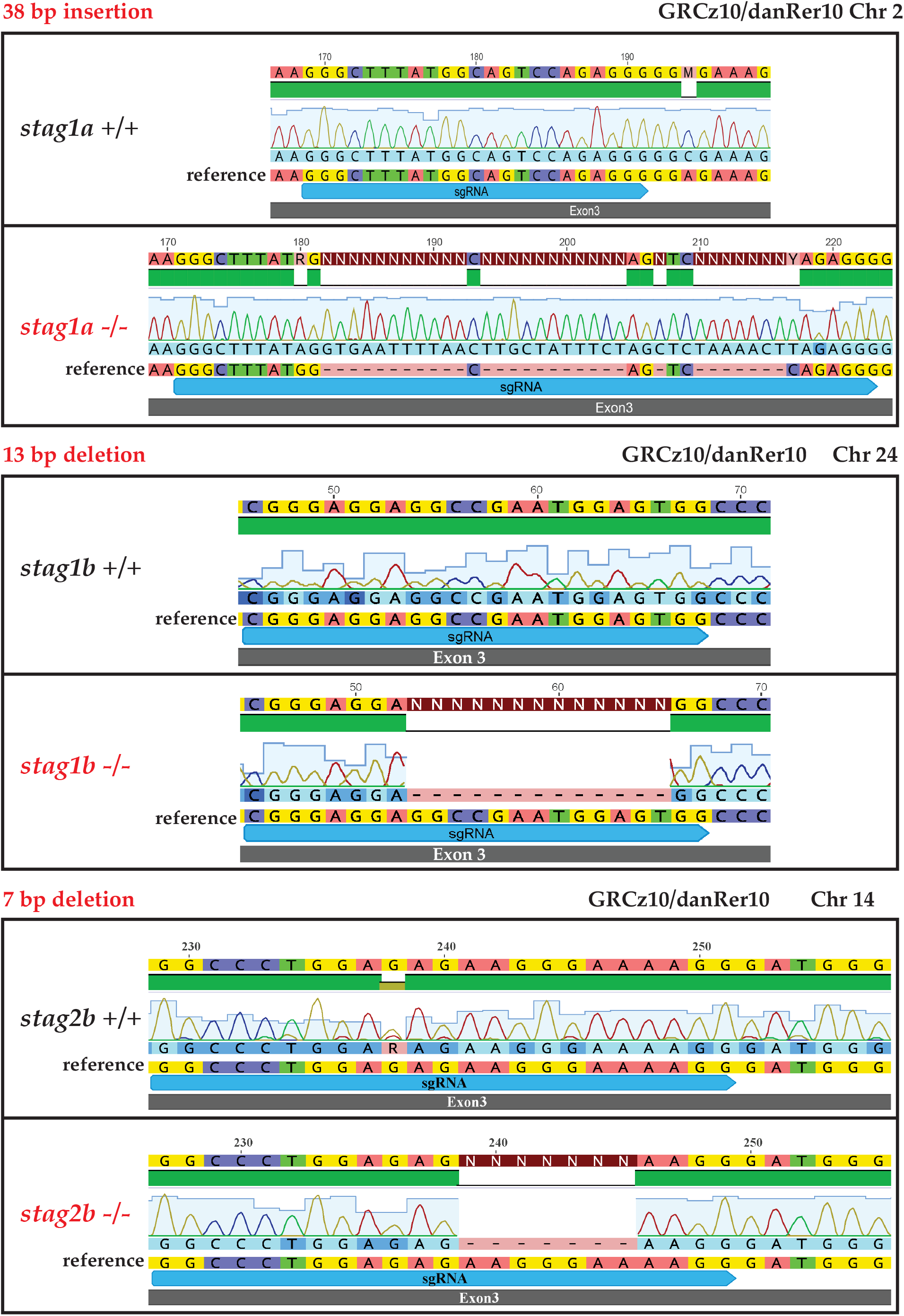
Genomic DNA sequence detail of zebrafish *stag* gene germ-line CRISPR mutants. Nucleotide alignments of wildtype and mutant homozygous sequences are shown. The sgRNA sites are annotated in blue.

**Supplementary Figure 3.**
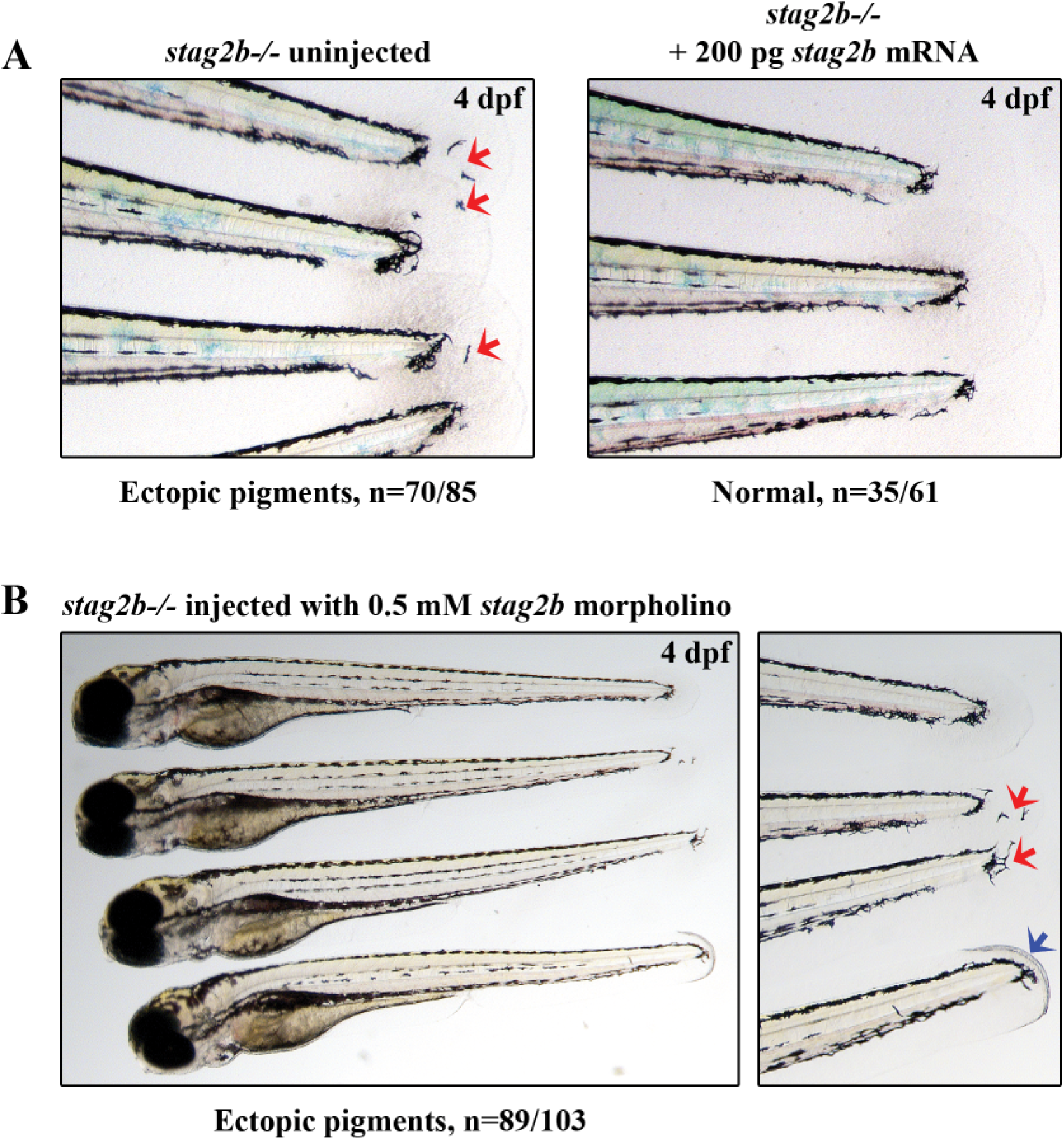
The *stag2b* mutation: confirmation of loss of function and and mutant phenotype. **(A)** The ectopic pigment cells seen in the tail fin of *stag2b*−/− embryos (red arrows) are rescued upon injection of *stag2b* mRNA. Lateral views of tail fin zoom-ins are shown, anterior to the left. **(B)** Injection of 0.5 mM *stag2b* morpholino does not induce any additional phenotypes in *stag2b*−/− embryos, and confirms that Stag2 loss causes displaced pigment cells (red arrows) and tail fin folds (blue arrow). Lateral views of full-length embryos and tail fin zoom-ins are shown, anterior to the left. Numbers of embryos are indicated below the respective panels.

**Supplementary Figure 4.**
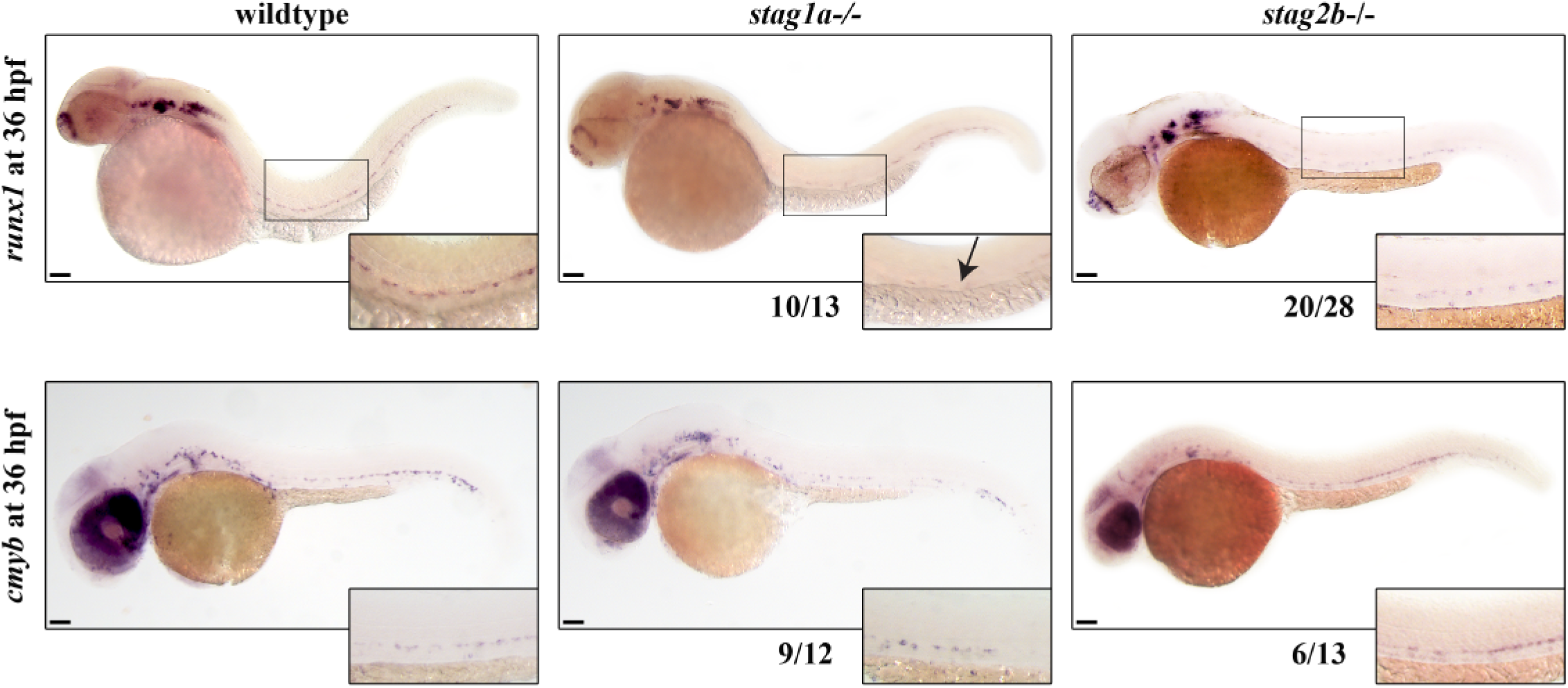
Whole-mount *in situ* hybridization analysis of expression of *runx1* and *cmyb* at 36 hpf. Lateral views are shown, anterior to the left. Insets show zoom-ins of the dorsal aorta region. Reduced *runx1* expression in *stag1a*−/− embryos is indicated by an arrow. Number of embryos is indicated below the respective panels. Scale bars are 100 *μ*m.

**Supplementary Figure 5.**
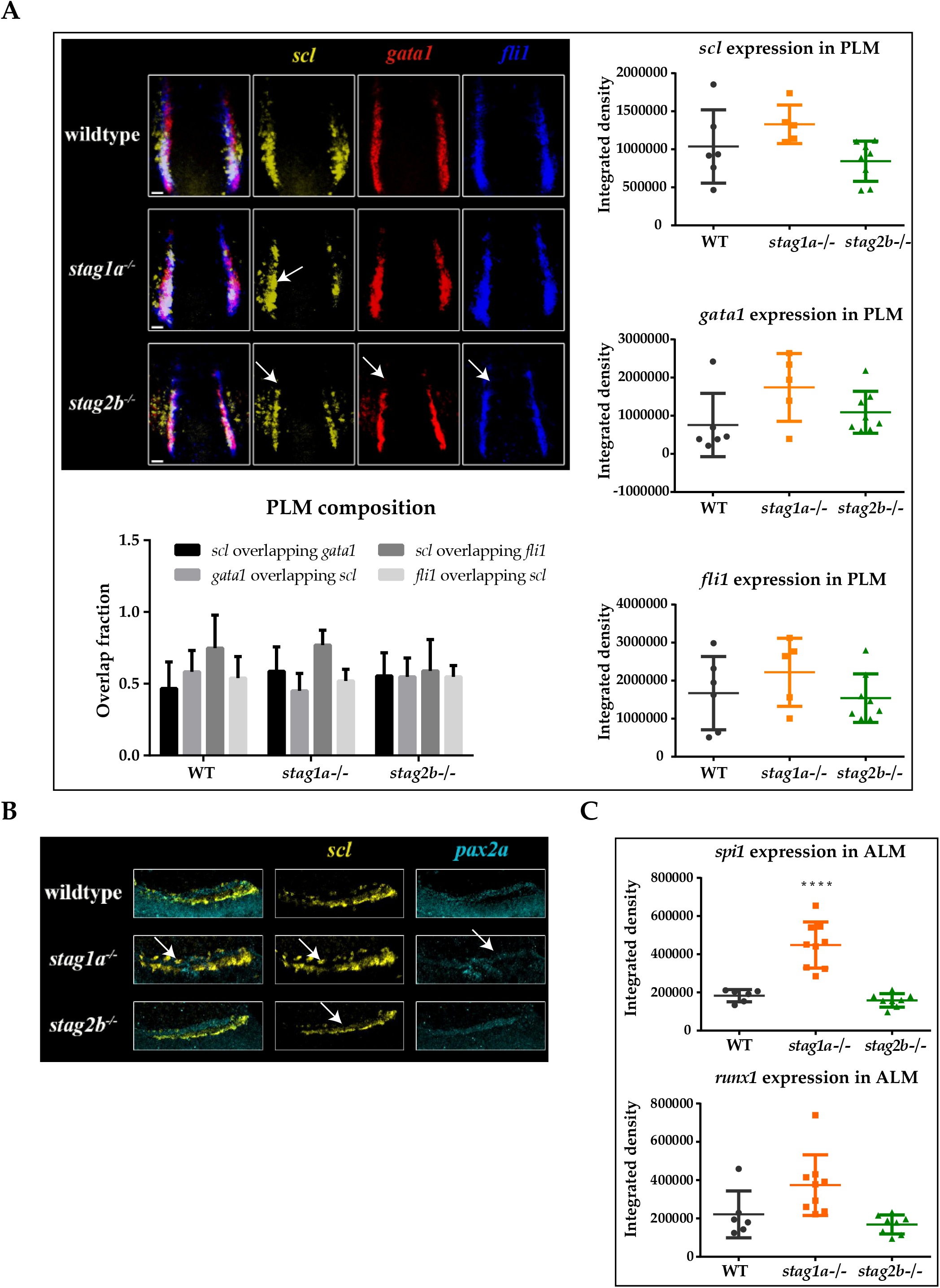
Stag mutations affect gene expression in the posterior lateral mesoderm (PLM) and anterior lateral mesoderm (ALM). **(A)** Multiplexed *in situ* HCR of *scl* (yellow), *gata1* (red), and *fli1* (blue) expression at 15 hpf. Dorsal PLM views are shown, anterior to the top. Extra *scl* expression in *stag1a*−/− embryos and reduction of PLM expression in *stag2b*−/− embryos is marked by white arrows. Quantitative analysis of fluorescence integrated densities indicates *scl*, *gata1* and *fli1* trend to non-significant upregulation in *stag1a*−/− embryos in the PLM. Composition of the PLM is equivalent in all embryos. **(B)** Multiplexed *in situ* HCR of *scl* (yellow) and *pax2a* (cyan) expression at 15 hpf. Posterior views of a single PLM stripe are shown, dorsal to the left. Arrows mark lateral expansion of *scl* into the *pax2a* domain in the middle *stag1a*−/− panels and reduced *scl* expression in the lower *stag2b*−/− panel. **(C)** Quantitative analysis of fluorescence integrated densities of ALM markers shows an increase of *runx1* and *spi1* expression in *stag1a*−/− embryos. **** P ≤ 0.0001; one-way ANOVA.

**Supplementary Figure 6.**
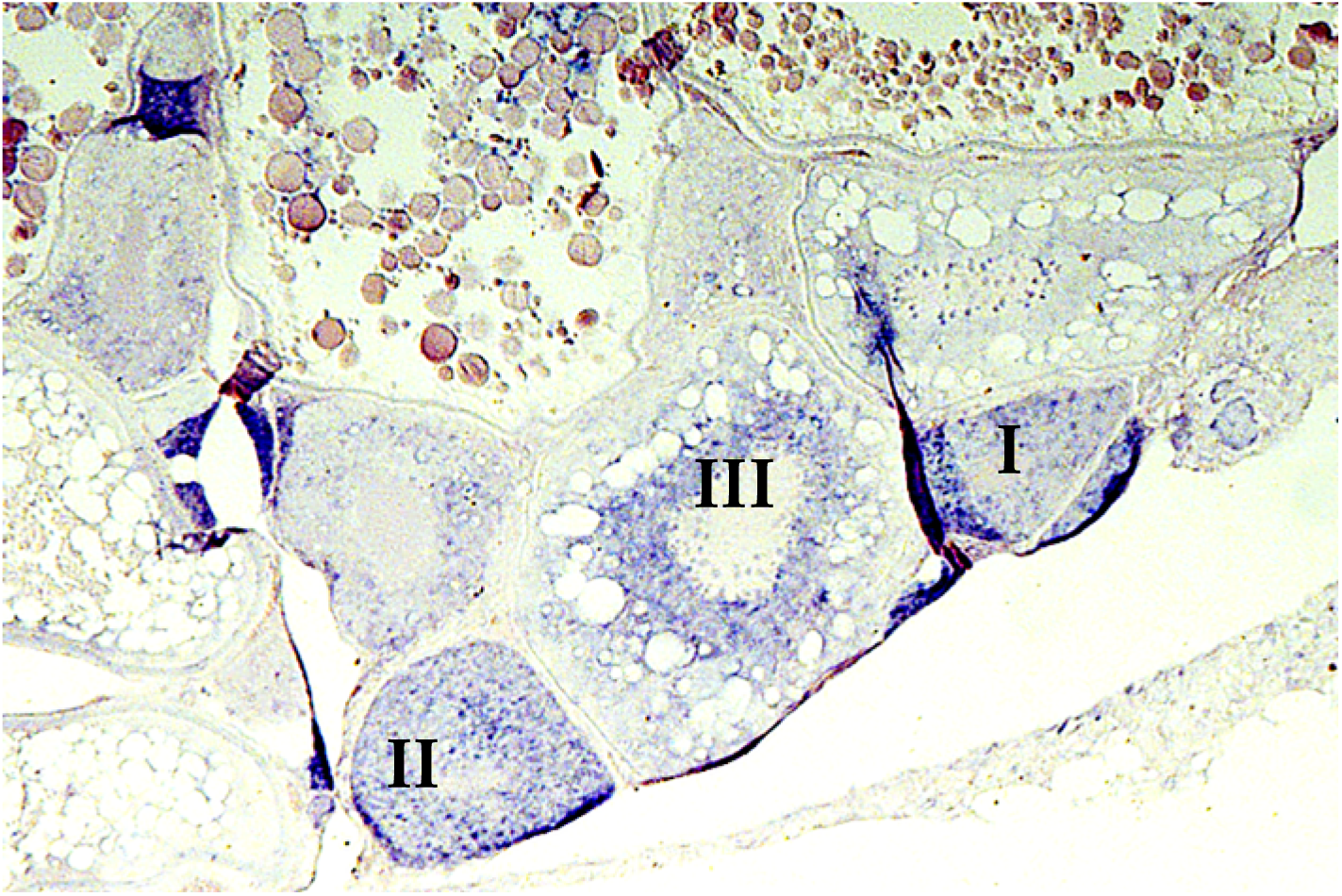
*in situ* hybridization of *stag2a* in adult zebrafish ovary. Expression of *stag2a* as detected by *in situ* hybridization (blue/purple) is clearly visible in Stage I, II and III oocytes in this transverse section through wild type adult zebrafish ovary.

**Supplementary Table 1.** List of accession identifiers for proteins used for phylogenetic analysis.

| Protein | Accession ID |
| --- | --- |
| Hs STAG1 | NP_005853.2 |
| Hs STAG2 | NP_001036214.1 |
| Gg STAG1 | XP_015146838.1 |
| Gg STAG2 | XP_004940885.1 |
| Mm Stag1 | NP_001344193.1 |
| Mm Stag2 | NP_001071180.1 |
| Xt stag1 | NP_001121432.1 |
| Xt stag2 | XP_002931833.2 |
| Dr stag1a | NP_001349269.1 |
| Dr stag1b | XP_692120.3 |
| Dr stag2a | NP_001093498.1 |
| Dr stag2b | XP_005173250.1 |

**Supplementary Table 2.**
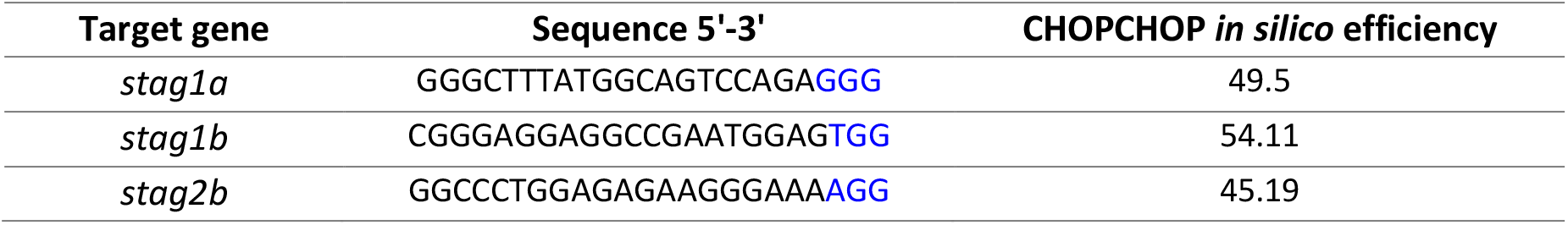
sgRNA sequences used to generate CRISPR mutants. PAM sequences are marked in blue.

**Supplementary Table 3.**
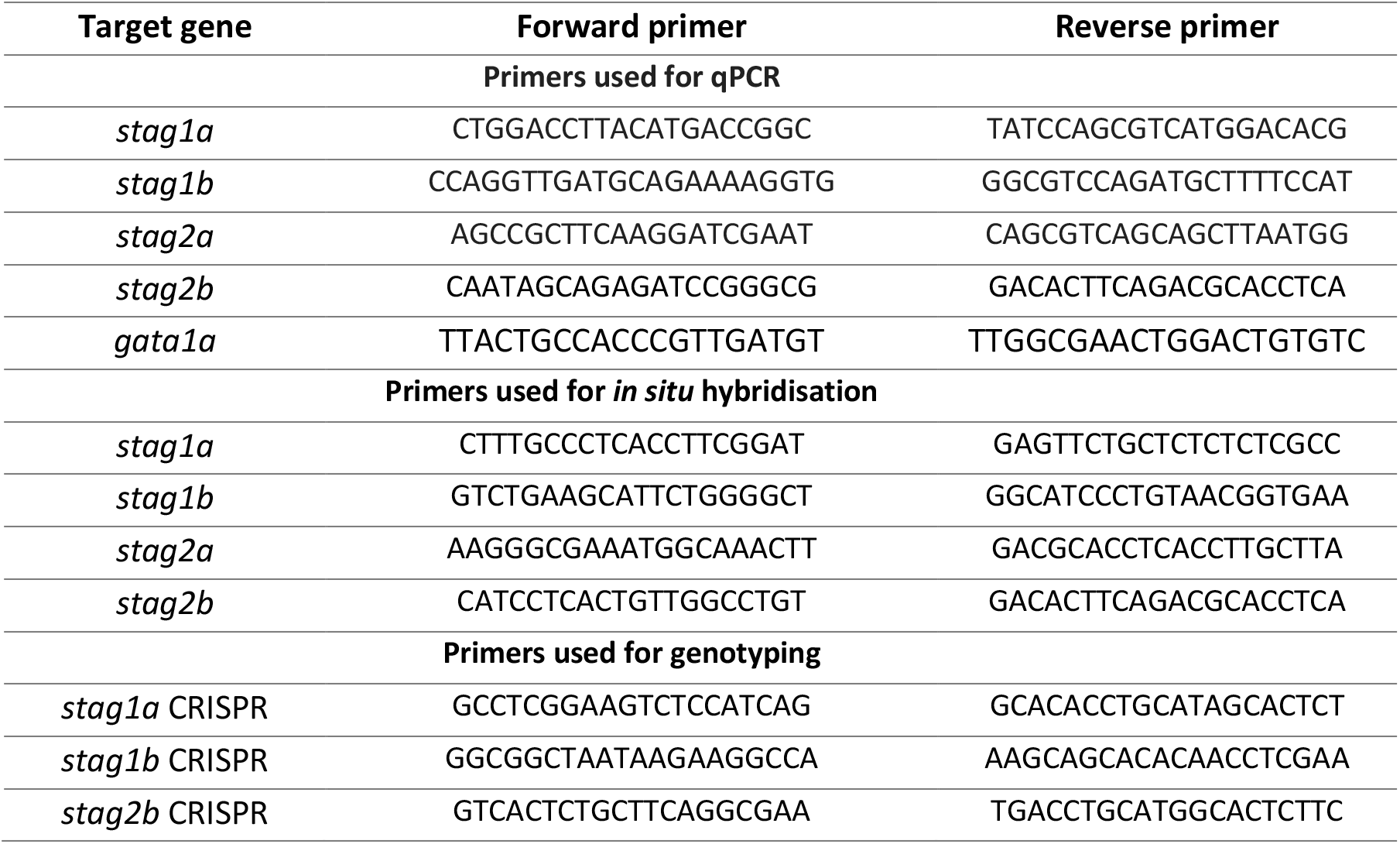
Primer sequences used in this study.

**Supplementary Table 4.**
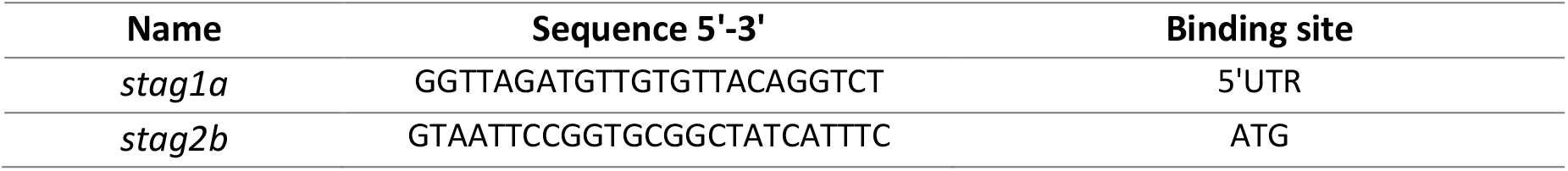
Morpholino sequences used in this study.

